# Protein Organization with Manifold Exploration and Spectral Clustering

**DOI:** 10.1101/2021.12.08.471858

**Authors:** Geoffroy Dubourg-Felonneau, Shahab Shams, Eyal Akiva, Lawrence Lee

**Affiliations:** Shiru, INC.; University of Texas at Dallas

## Abstract

We present a method to provide a biologically meaningful representation of the space of protein sequences. While billions of protein sequences are available, organizing this vast amount of information into functional categories is daunting, time-consuming and incomplete. We present our unsupervised approach that combines Transformer protein language models, UMAP graphs, and spectral clustering to create meaningful clusters in the protein spaces. To demonstrate the meaningfulness of the clusters, we show that they preserve most of the signal present in a dataset of manually curated enzyme protein families.

## 1 Introduction

Current gene sequencing and structure prediction technologies generate an ever-increasing number of protein sequences and their respective highly confident structural models [1]. This exponential growth [2] in data availability is contrasted by the slow rate of functional characterization of proteins. This lag between sequences and their respective functions stems from the requirement of multiple years of meticulous work to elucidate the many aspects of functions of even a single protein. Automating efficient function prediction is thus a “holy grail” that can transform multiple fields of academic and applied research. Our hope is that such a method clusters protein sequences to into iso-functional groups in different levels of granularity. Several “gold standard” hierarchical databases that map protein sequence to folds, domains and biochemical functions of proteins can be used for validating such function prediction algorithms, for example the Protein Structure Classification Database (CATH)[3], Protein Families (Pfam)[4] and the Structure Function Linkage Database (SFLD)[5]. These databases are drawn from curating the scientific literature coupled with the careful development of models for protein categorization, mainly dependent upon percent sequence identity, or statistically significant hits of the sequence patterns, i.e. Hidden Markov Models). Developing methods that achieve equal or more accurate function prediction can bridge the ever-increasing gap between sequence abundance and sparse functional data.

In this paper, we introduce a computational method that is capable of providing competitive results of protein function in intervals of a few minutes. Our method enables scientists to propagate the grouping knowledge to niches of protein sequence space that lack any functional annotation, as well as to adjust the grouping granularity for each purpose (which is not always an option when using a pre-compiled database).

In the pre-processing, we embed sequence data using facebook/esm-1b [6]. This allows us to benefit from fixed-size numerical feature vectors. The first task is to find a suitable similarity measure, Learning Meaningful Representations of Life (NeurIPS 2021), Virtual conference. denoted as 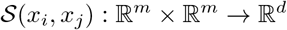 where *m* is the dimension of the embedding space, and *d* is the dimension of similarity operator. Here, for the sake of simplicity, we set *d* = 1. Then, based on the obtained 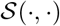, we proceed by clustering proteins into clusters that preserve functionalities.

Since we define our challenge as an unsupervised learning problem, we evaluate the precision of our proposed method by comparing the clusters that our algorithm yields with popular sequence to function assignments based on the experimental-based sequence-function mappings by the SFLD database.

## 2 Method

The outline of this work is divided into two major sections. The first subtask is searching for a novel similarity measure between proteins. The second subtask is clustering data points.

### 2.1 Manifold Representations

The embedded sequences are conventionally presented in the standard Riemannian structure consisting of the Euclidean space equipped with the Euclidean metric. However, in the standard system, similar proteins do not necessarily have similar functionalities, and proteins that are not similar in sequence might be similar in functionalities. This inconsistency provokes the question of whether this linear system suits the task of protein clustering. To address this issue, we generalize the standard Riemannian manifold by substituting the Euclidean space with a nonlinear but smooth manifold. The condition on the Riemannian metric can also be relaxed by allowing multiple metrics that come from a diverse set and are specific to each data point. The goal is to find a nonlinear system whose notion of similarity reliably translates into similarity in function. Uniform Manifold Approximation and Projection (UMAP) [7] provides a general platform within which we can approximate the Riemannian manifold we assume the data lies on. There are other methods that serve the same purpose, such as t-SNE [8], but UMAP is arguably more advantageous in preserving the global structure of the data.

UMAP results in a manifold that can be described as a weighted graph, *G* = (*V_G_, E_G_, W_G_*), such that *V_G_* is the set of all proteins, and for each (*i*, *j*) ∈ *E_G_, W_i,j_* determines the magnitude of similarity between *x_i_* and *x_j_*, which is equivalent to 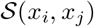. This approach enables us to assign a unique similarity measure to each protein. And each 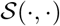 is flexible to be learned or set based on expert’s opinion or heuristically. In addition, UMAP provides us with 2-dimensional visualizations. The next step is to investigate whether the nonlinear system admits any advantage for clustering.

### 2.2 Node Clustering

The manifold obtained from UMAP is a weighted graph. Thus, a natural proceeding would be to perform node clustering. However, since this is an unsupervised task, there is no reliable way to determine the number of clusters. Even worse, identifying communities is an ill-defined problem, meaning that there is no clear definition of community, nor a definite criterion for assigning nodes to clusters [9]. In our particular example, each protein might have several functionalities. Hence, there might be multiple suitable clusters for each protein. As a result, the objective function of any protein clustering method should be exclusively designed based on the particular set of functionalities that one might want to find a substitution for.

The absence of a clear definition of community causes an incapacitating uncertainty in the search for benchmarks. To have a guideline that is acceptable in the field, given that complex graphs are usually hierarchical in essence, we suggest following the style of previously constructed databases, such as SFLD and CATH, and categorize proteins into a hierarchy of clusters. Hierarchy has multiple advantages such as flexible granularity and fewer hyperparameters. Since actual clusters might not be balanced or evenly distributed, hierarchy enables us to have the option of balanced methods, and still get the clustering that describes the data set accurately. In addition, taking any established categorization as a benchmark can cut the expenses of evaluation that otherwise would have to be done via highly expensive, time-consuming, and almost impossible procedures. We set our objective to implement a downsampling operation that at each round gathers sequences with the same functionalities within the same cluster while getting lower variance in functionalities in each cluster. Learning the hierarchy of complex graphs has been done via a variety of methods [10]. Since Graclus [11], methods like DiffPool [12] [13] learn graph hierarchical structures based on pooling modules. Of our special interests are methods that take the graph topology and the node features jointly, such as EigenPooling [14] that extracts subgraph information via local graph Fourier transform, or SAGPool [15] that uses self-attention. Recently, in minCutPool [16] authors take the general technique of spectral clustering, and perform an efficient pooling to generate a coarsened version of the graph. However, Tsitsulin et al.[17] shows some inaccuracies in minCutPool optimization and proposes an alternative, modularity-based technique that outperforms the aforementioned methods. Intuitively, spectral modularity objective [18] quantifies the quality with respect to a random graph.

Escaping local minimas is an important challenge in unsupervised node clustering. However, Good et al. [19], show that with the modularity objective function, graphs with hierarchical structure exhibit many degenerate solutions, but the modularity scores of alternative solutions can be very close to the global optima. Our experiments with the UMAP graph confirm both claims. Modularity might end up in a suboptima, but all the observed local minimas admit acceptable results. In addition, modularity objective function works well when nodes have the same degree distribution, which also holds for our case. Hence, we pick Deep Modularity Networks (DMoN) [17] to proceed. Here we skip representation of DMoN, but encourage you to check their detailed explanation.

Our approach is similar to the downsampling done by Simonovsky et al. [20], where the authors divided the data points into two clusters. Our branching factor is three because we have not seen any clear advantage from two, and indeed three reaches the practical granularity faster.

## 3 Results

### 3.1 Database

We chose to evaluate our method using a dataset of enzymes. An enzyme can be assigned with what is considered as one of the most accurately defined “protein functions”. The SFLD is a hierarchical classification system of enzymes that maps protein sequence to specific chemical reactions. At the highest level, the SFLD categorizes enzymes into 11 functionally diverse enzyme superfamilies. A superfamily is a broad set of evolutionarily related enzymes with a shared chemical function that maps to a conserved set of active site features. Members of the same enzyme superfamilies can share very little sequence similarity, but catalyze the same chemical reaction, or vice-versa: they can share high level of sequence similarity, but perform diverse chemical reactions. Each superfamily is further divided into subgroups. A subgroup is a set of evolutionarily-related enzymes that have more shared features among each other relative to other members of the superfamily; they may still catalyze different chemical reactions. At the most granular level, each subgroup consists of several families. A family is a set of evolutionarily-related enzymes that catalyze the same chemical reaction.

We define all members of an enzyme family as having the same function. Hence, recapitulating SFLD families using our unsupervised method is an indication that our method is actually capable of predicting functionalities.

### 3.2 Discussion

Members of the same enzyme family were assigned manually by experts, via literature-based documentation [5]. As such, they are very likely to have the same functionalities. Hence, we evaluate our final clustering by quantifying how well each family has been preserved in the obtained clusters. Preservation percentage is defined as the ratio of the largest division of a family to the the total size of the family. For example, having 3 clusters, if 70% of a family is clustered into one cluster, and the remaining 30% is clustered into another cluster, the preservation is 70%.

Finding a practical replacement for a protein can be done by gradually narrowing down the options, until reaching a set of proteins small enough for further investigations. The idea is not to eliminate any protein that can effectively be a good substitution in the process. The main challenge is to keep proteins that are not very similar in their sequence, but share the same functionalities.

This result might not be flawless, but it shows a considerable potential for further investigation. For further enhancements, one can expand the initial feature vectors, or derive similarity measures based on experts’ opinion. Having a modularity objective function, it is convenient to think of a community as a subset of nodes such that the probability of nodes being connected to inside nodes is higher than outside nodes. Approximating these probabilities could be a well-defined problem if there was a model stating how edges are formed. This is a possible place where experts’ knowledge can be incorporated (e.g. using phylogenetic information)

## 4 Conclusion

The method presented here can help researchers assign and test functional properties of proteins from the vast niches of functionally unexplored sequence space. Such predictions could dramatically shorten the time of experimental validation of novel protein functions. Furthermore, through the use of protein embeddings it allows one to capture similarities beyond the traditional means (e.g., percent identity). As such, this method can be used to identify proteins with improved function (e.g., thermal stability and yield) for applied science and protein engineering.

**Figure 1:**
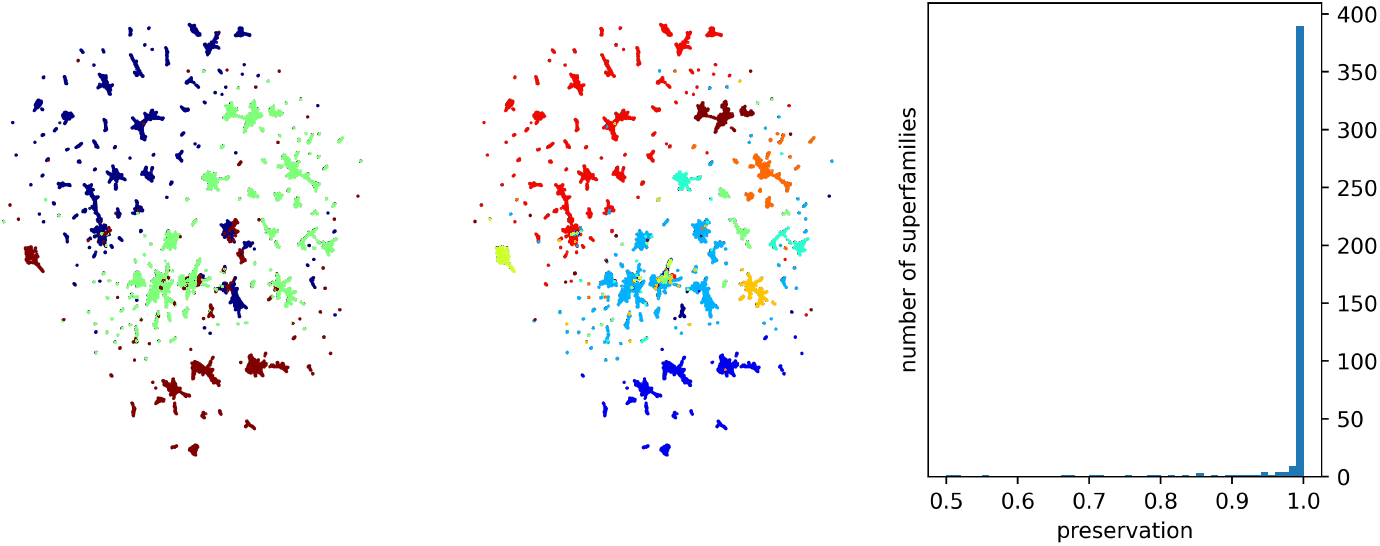
The first round of manifold projection and unsupervised clustering over SFLD. The figures show the 2-D presentation of 332,090 enzymes from the 11 superfamilies of SFLD. This 2-D presentation is obtained by UMAP with *n_neighbors* = 250, *random_state* = 42, *min_dist* = 0, and *metric* = Euclidean. In the middle plot, each color identifies a superfamily. The left figure is the result of clustering SFLD into 3 clusters with DMoN. SFLD consists of 440 families, including families of enzymes within each subgroup that have not been assigned to a family, as well as families of enzymes within each superfamily without subgroup assignment. The histogram shows the first round of unsupervised clustering preserves 390 families 99% and higher.

**Figure 2:**
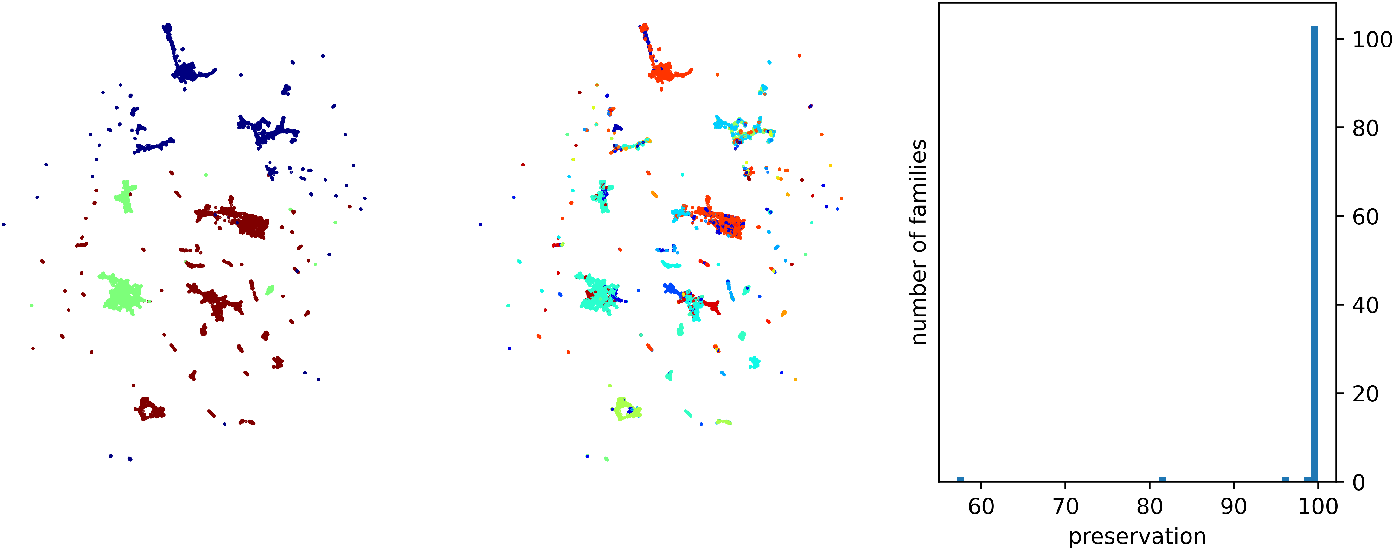
The second round of manifold projection and unsupervised clustering over one of the clusters obtained from the first round. This cluster consists of 69,484 enzymes, including 99.8% of the superfamily Enloase, and 98.6% of the superfamily Radical SAM Phosphomethylpyrimidine Synthase. In the middle plot, each color identifies a family. Out of 107 families, our clustering preserves 103 clusters higher than 99%. In the left plot, each color is a cluster.

